# Exploring the causal effects of genetic liability to ADHD and Autism on Alzheimer’s disease

**DOI:** 10.1101/2020.04.15.043380

**Authors:** Panagiota Pagoni, Christina Dardani, Beate Leppert, Roxanna Korologou-Linden, George Davey Smith, Laura D Howe, Emma L Anderson, Evie Stergiakouli

## Abstract

**Background:** There are very few studies investigating possible links between Attention Deficit Hyperactivity Disorder (ADHD), Autism Spectrum Disorder (ASD) and Alzheimer’s disease and these have been limited by small sample sizes, diagnostic and recall bias. However, neurocognitive deficits affecting educational attainment in individuals with ADHD could be risk factors for Alzheimer’s later in life while hyper plasticity of the brain in ASD and strong positive genetic correlations of ASD with IQ and educational attainment could be protective against Alzheimer’s.

**Methods:** We estimated the bidirectional total causal effects of genetic liability to ADHD and ASD on Alzheimer’s disease through two-sample Mendelian randomization. We investigated their direct effects, independent of educational attainment and IQ, through Multivariable Mendelian randomization.

**Results:** There was limited evidence to suggest that genetic liability to ADHD (OR=1.00, 95% CI: 0.98 to 1.02, p=0.39) or ASD (OR=0.99, 95% CI: 0.97 to 1.01, p=0.70) was associated with risk of Alzheimer’s disease. Similar causal effect estimates were identified when the direct effects, independent of educational attainment (ADHD: OR=1.00, 95% CI: 0.99 to 1.01, p=0.07; ASD: OR=0.99, 95% CI: 0.98 to 1.00, p=0.28) and IQ (ADHD: OR=1.00, 95% CI: 0.99 to 1.02. p=0.29; ASD: OR=0.99, 95% CI: 0.98 to 1.01, p=0.99), were assessed. Finally, genetic liability to Alzheimer’s disease was not found to have a causal effect on risk of ADHD or ASD (ADHD: OR=1.12, 95% CI: 0.86 to 1.44, p=0.37; ASD: OR=1.19, 95% CI: 0.94 to 1.51, p=0.14).

**Conclusions:** In the first study to date investigating the causal associations between genetic liability to ADHD, ASD and Alzheimer’s, within an MR framework, we found limited evidence to suggest a causal effect. It is important to encourage future research using ADHD and ASD specific subtype data, as well as longitudinal data in order to further elucidate any associations between these conditions.

## INTRODUCTION

Attention Deficit Hyperactivity Disorder (ADHD) and Autism Spectrum disorder (ASD) are lifelong neurodevelopmental conditions associated with large societal costs (1-3). It has been estimated that the largest proportion of these costs is attributed to the increased physical care and psychosocial support needs of the affected individuals during adulthood (4, 5). Despite increasing interest on the adult outcomes of ADHD and ASD, there is currently limited evidence on the associations of both neurodevelopmental conditions with one of the most debilitating conditions of old age, Alzheimer’s disease. Research in this area is of great importance in order to inform family support and financial planning as well as societal policies and services.

ADHD is characterised by difficulties in several areas of neurocognitive functioning, including memory, attention and inhibitory control (6). In later life, these difficulties could be risk factors for an Alzheimer’s disease diagnosis (7, 8). This relationship might be mediated by educational attainment and IQ (9). ADHD has been associated with lower educational attainment and IQ in observational and genetic studies (9, 10), and they in turn confer increased risk for Alzheimer’s (11, 12).

In the case of ASD, it has been suggested that abnormal synaptogenesis, connectivity and hyper plasticity of the brain identified recently in ASD, could be protective against Alzheimer’s disease (13). Adding to this, a protective effect of ASD on risk of Alzheimer’s could be further hypothesized based on evidence suggesting strong positive genetic correlations of ASD with IQ and educational attainment (14), which could therefore mediate any associations between the two conditions.

Beyond these hypotheses, little is currently known on the possible associations between ADHD, ASD and Alzheimer’s. There is only a small number of case-control studies on ADHD, suggesting a higher frequency of ADHD symptoms in patients with dementia with Lewy body and a higher risk of dementia in adults with ADHD (15-17). However, these studies are limited by small sample sizes and the possibility of diagnostic and recall bias, since sample definition has mostly relied on self or informant reports (18). In addition, longitudinal studies require a long follow-up period (from childhood to late adulthood) and could be biased by attrition and confounding. Finally, despite the availability of genome-wide association study (GWAS) data on ADHD, ASD and Alzheimer’s disease (14, 19, 20) studies have focused solely on the genetic correlation between the three phenotypes, showing limited evidence of a genetic overlap (14, 19, 20).

The release of the latest ADHD and ASD GWASs provides a unique opportunity to investigate the possible causal associations between genetic liability to ADHD, ASD and Alzheimer’s, through a method that overcomes limitations of observational studies; Mendelian randomization. Mendelian randomization is an instrumental variables (IVs) approach using genetic variants as proxies for exposures to investigate the causal effects of these exposures on health outcomes (21). As genetic variants are randomly allocated at meiosis and fixed at conception, the method is robust to confounding and reverse causation (22).

For these reasons, we aimed to investigate the bidirectional causal associations between genetic liability to ADHD, ASD and Alzheimer’s using two-sample Mendelian randomization. Next, we performed Multivariable Mendelian randomization to estimate the direct causal effects of genetic liability to ADHD and ASD on risk of Alzheimer’s, independent of educational attainment and IQ to explore the possibility that the causal effect of ADHD and ASD on Alzheimer’s are being masked by IQ or educational attainment.

## METHODS

### Two-sample Mendelian randomization (MR)

MR relies on strict assumptions that the genetic variants should satisfy in order to be considered valid instruments and therefore yield unbiased causal effect estimates. Specifically, the genetic variants:

A. must be strongly associated with the exposure,
B. independently of any confounders of the exposure-outcome association, and
C. are associated with the outcome only via their effect on the exposure (i.e. absence of horizontal pleiotropy).

In the context of the present study we applied two-sample MR, a method that enhances statistical power and the precision of the causal effect estimates. This is because the method does not require data on exposure, outcome and genotype in a single sample. Instead, instrument-exposure and instrument-outcome effect sizes and standard errors are extracted from GWAS conducted in independent samples of the same underlying population.

### GWAS Summary Data

We used GWAS summary data from the latest publicly available GWASs on the exposures of interest. Table 1 provides a summary of the studies utilized. Detailed information can be found in the original publications (14, 19, 20, 23, 24).

**Table 1.** Summary of the GWASs used in two-sample Mendelian randomization analyses & multivariable Mendelian randomization. Samples in bold indicate overlap between the different GWASs used.

| Phenotype | PMID | Cohorts & samples | N |
| --- | --- | --- | --- |
| ADHD | Demontis et al.<br>(2018)<br>PMID: 30478444 | iPSYCH & PGC consortium | 20,183 cases<br>35,191 controls |
| ASD | Grove et al.<br>(2019)<br>PMID: 30804558 | iPSYCH & PGC consortium | 18,381 cases<br>27,969 controls |
| Alzheimer's | Jansen et al.<br>(2019)<br>PMID: 30617256 | Phase 1:<br>PGC-ALZ (DemGene, **TwinGene**, STSA),<br>IGAP, ADSP<br>Phase 2:<br>Proxy-cases of Alzheimer's disease in **UKB**<br>Phase 3:<br>Meta-analysis of Phase 1&2 | Phase1:<br>24,087 cases<br>55,058 controls<br>Phase 3:<br>71,880 cases<br>383,378 controls |
| Educational<br>Attainment<br>(Years of<br>Schooling) | Lee et al<br>(2018)<br>PMID: 30038396 | Add Health, EGCUT , ELSA, FENLAND,<br>Geisinger, GSII, NORFOLK, **UKB**, UKHLS,<br>VIKING, WLS,ACPRC, AGE, ALSPAC,<br>ASPS, BASE-II,CoLaus,COPSAC2000,<br>CROATIA-Korčula, deCODE, DHS, DIL,<br>ERF, FamHS, FINRISK, FTC, GOYA,<br>GRAPHIC, GS, H2000 Cases, H2000 Controls,<br>HBCS, HCS, HNRS (CorexB), HNRS (Oexpr),<br>HNRS (Omni1), HRS, Hypergenes, INGI-<br>CARL, INGI-FVG, KORA S3, KORA S4,<br>LBC1921, LBC1936, LifeLines, MCTFR,<br>MGS, MoBa, NBS, NESDA, NFBC66 , NTR,<br>OGP, OGP-Talana, ORCADES, PREVEND,<br>QIMR, RS-I, RS-II, RS-III, Rush-MAP, Rush-ROS, SardiNIA, SHIP, SHIP-TREND, STR –<br>Salty, STR – **Twingene**, THISEAS, TwinsUK,<br>WTCCC58C, YFS | 766,345<br>participants |
| Intelligence<br>(IQ) | Savage et al<br>(2018)<br>PMID: 29942086 | **UKB**, COGENT, RS, GENR, STR, S4S,<br>HiQ/HRS, TEDS, DTR-MADT, DTR-LSADT,<br>IMAGEN, BLTS-Children, BLTS-Adolescence, NESCOG, GfG, STSA-<br>SATSA+GENDER, STSA-HAMRONY | 269,867<br>participants |

#### Instrument extraction

For each exposure of interest independent genetic variants were identified (r^2^<0.01 within a 10,000 kb window, p<5×10^−08^) and corresponding log odds ratios and standard errors were extracted from the publicly available datasets. An exception was ASD, where only two independent variants were identified. Therefore, in order to increase the power of our study, we relaxed the significance threshold (p <5×10^−07^) for instrument selection. A similar approach for ASD instrument selection has been used in a previous study (25). Exposure instruments were extracted from the outcome GWAS. When a genetic variant was not present in the outcome GWAS, we identified proxy variants using the LDLink online tool (26) (r^2^<0.01 within a 10,000 kb window).

#### Harmonisation

As instrument-exposure effect estimates were coded to express the effect estimate per increasing allele, the alleles of the variants identified in the outcome were harmonised so that their effect estimates corresponded to the alleles of the exposure. The GWAS summary data of ADHD and ASD do not offer information on effect allele frequencies. Therefore, for variants where the alignment of the alleles between the exposure and outcome variants was not possible, were excluded as palindromic. Information on the genetic variants included in the analysis can be found in Supplementary material (Table S1-S3). The analytic process that was followed across the analyses of the present study is illustrated in FigureS1.

### Statistical Analyses

Causal effect estimates were generated using the Inverse-Variance-Weighted (IVW) method. IVW can be used to summarize the causal effects of multiple genetic variants, as it is equivalent to fitting a weighted linear regression of the gene-outcome associations on the gene-exposure associations, with the intercept term constrained to zero (27, 28). Thus, IVW estimates assume that all genetic variants are valid instruments with no pleiotropic effects.

### Sensitivity Analyses

In order to explore the validity of MR assumptions, we compared the estimated total causal effects obtained from univariable MR to the effects obtained using the MR-Egger regression and the Weight median estimator (29, 30). Briefly, MR-Egger regression unlike the IVW method, allows for an unconstrained intercept term and therefore the intercept term is a formal statistical test for the presence of horizontal pleiotropy, while the slope provides a causal effect estimate accounting for pleiotropic bias. The Weighted median estimator provides a causal effect estimate even when up to half of the genetic variants are invalid instruments, by estimating the causal effect as the median of the weighted ratio estimates. Consistent results across the above methods are indicative of a true causal effect.

Additionally, as the validity of MR results depends largely on the strength of the genetic instruments, we used F-statistic to ensure that weak instrument bias will not affect our results (31). F-statistic estimates the instrument strength of a genetic variant, which is a function of the variance explained by a set of genetic variants (R^2^), the number of genetic variants used and the sample size. An F-statistic smaller than 10 indicates the presence of weak instrument bias and that causal effect estimates are likely to be influenced.

The influence of each genetic variant on the outcome, was assessed via a leave-one out analysis, in which genetic variants are systematically removed and causal effects of the remaining SNPs on the outcome are re-estimated (32). Finally, the Cochran’s Q statistic was used, to assess whether the causal estimates of the genetic variants were comparable (30). Observed substantial heterogeneity was used as an indication that genetic variants may not be valid instruments.

### Multivariable MR

MVMR is an extension of MR that can be utilized in cases that multiple exposures seem to be strongly genetically related and are considered to have possible causal effects on an outcome. MVMR allows the estimation of the direct effects of each exposure on the outcome by entering the exposures within the same model. Detailed information on the method can be found in the original publication (33).

### Analyses restricted to clinically diagnosed Alzheimer’s disease cases

The main analyses conducted in the present study used summary data from Phase 3 of the Alzheimer’s GWAS, which included proxy-cases of Alzheimer’s disease in UK Biobank (Table1). This might be problematic as: i. participants were defined as cases, based on family history, ii. which was self-reported, iii. and they were asked about broad dementia and not Alzheimer’s specifically. In addition, UK Biobank is an overlapping sample across the Alzheimer’s, educational attainment and intelligence GWASs and this might bias current analyses (34). More specifically, the overlap of participants between the Alzheimer’s and educational attainment GWAS was 58% and 72% with IQ GWAS.

Considering the above limitations, we conducted a second round of analyses using summary data of the meta-analysis of Korologou-Linden et al. (35) which corresponds to Phase 1 of the Alzheimer’s GWAS. Thus, it included only clinically diagnosed cases and the sample overlap between the Alzheimer’s GWAS and the educational attainment GWAS reduced to 1%.

## RESULTS

### Instrument Strength

Instrument strength as estimated by the F-statistic did not indicate weak instrument bias as it ranged from 30 to 51 for ADHD, 25 to 35 for ASD and 30 to 945 for Alzheimer’s instruments.

### Causal effects of genetic liability to ADHD on Alzheimer’s

#### i. Bidirectional total causal effects

A total of 10 genetic variants were used to estimate the total causal effect of genetic liability to ADHD on Alzheimer’s disease. We found limited evidence of a causal effect of genetic liability to ADHD (OR=1.00, 95% CI: 0.98 to 1.02, p=0.39) on Alzheimer’s disease. Horizontal pleiotropy was unlikely to bias the result, as suggested by the MR-Egger intercept term (OR=0.99, 95% CI: 0.99 to 1.00, p=0.91). Moreover, MR-Egger and weighted median estimators yielded similar results to the IVW estimator for ADHD (Table 2). No substantial heterogeneity was observed between the genetic variants, as indicated by Cochrane’s Q statistic (Q=6.52, p=0.68), and leave-one out analyses did not identify any SNP as influential (Figure S2A).

**Table 2.** Bidirectional effects of genetic liability to ADHD on Alzheimer’s disease

|  |  | Causal effect estimates |  |  |  |
| --- | --- | --- | --- | --- | --- |
|  | No. SNPs | OR | 95%CI | P-value | Q p-value |
| ADHD on Alzheimer's disease |  |  |  |  |  |
| IVW | 10 | 1.00 | 0.98, 1.02 | 0.39 | 0.68 |
| MR-Egger |  |  |  |  |  |
| Intercept | 10 | 0.99 | 0.99, 1.00 | 0.91 | - |
| Slope | 10 | 1.01 | 0.92, 1.11 | 0.76 | 0.58 |
| Weighted Median | 10 | 1.01 | 0.99,1.04 | 0.17 | - |
| Alzheimer's disease on ADHD |  |  |  |  |  |
| IVW | 31 | 1.12 | 0.86, 1.44 | 0.37 | 0.10 |
| MR-Egger |  |  |  |  |  |
| Intercept | 31 | 0.99 | 0.98, 1.01 | 0.97 | - |
| Slope | 31 | 1.12 | 0.77, 1.64 | 0.53 | 0.86 |
| Weighted Median | 31 | 1.12 | 0.81,1.56 | 0.47 | - |

IVW: Inverse-variance weighted; SNP – single nucleotide polymorphism; OR: Odds Ratio; CI: Confidence Intervals. Results are presented per log odds ratio increase of the exposure of interest.

In the reverse direction, 31 genetic variants were available to be used for estimation of the total causal effect. The IVW estimator suggested limited evidence of a causal effect of genetic liability to Alzheimer’s on ADHD (OR=1.12, 95% CI: 0.86 to 1.44, p=0.37). MR-Egger and Weight median estimators were directionally consistent with the IVW (Table 2). Heterogeneity among the variants was unlikely to bias the results (Q=39.84, p=0.10), and leave-one out analysis did not point out any genetic variant as influential (Figure S2B).

#### ii. Direct effects

Following the exclusion of all palindromic SNPs during harmonization and after clumping the final set of genetic variants (to ensure that only independent variants are included in our analysis), a total of 643 and of 180 variants were available for inclusion in the ADHD-EA-AD and ADHD-IQ-AD multivariable analyses respectively.

The estimate of the direct causal effect of genetic liability to ADHD on Alzheimer’s, independent of educational attainment, remained virtually the same (OR=1.00, 95% CI: 0.99 to 1.01, p=0.76) (Table 3A). A similar causal effect was identified after estimating the direct causal effect of genetic liability to ADHD on Alzheimer’s, independent of IQ (OR=1.00, 95%CI: 0.99 to 1.02. p=0.29) (Table 3B).

**Table 3.**
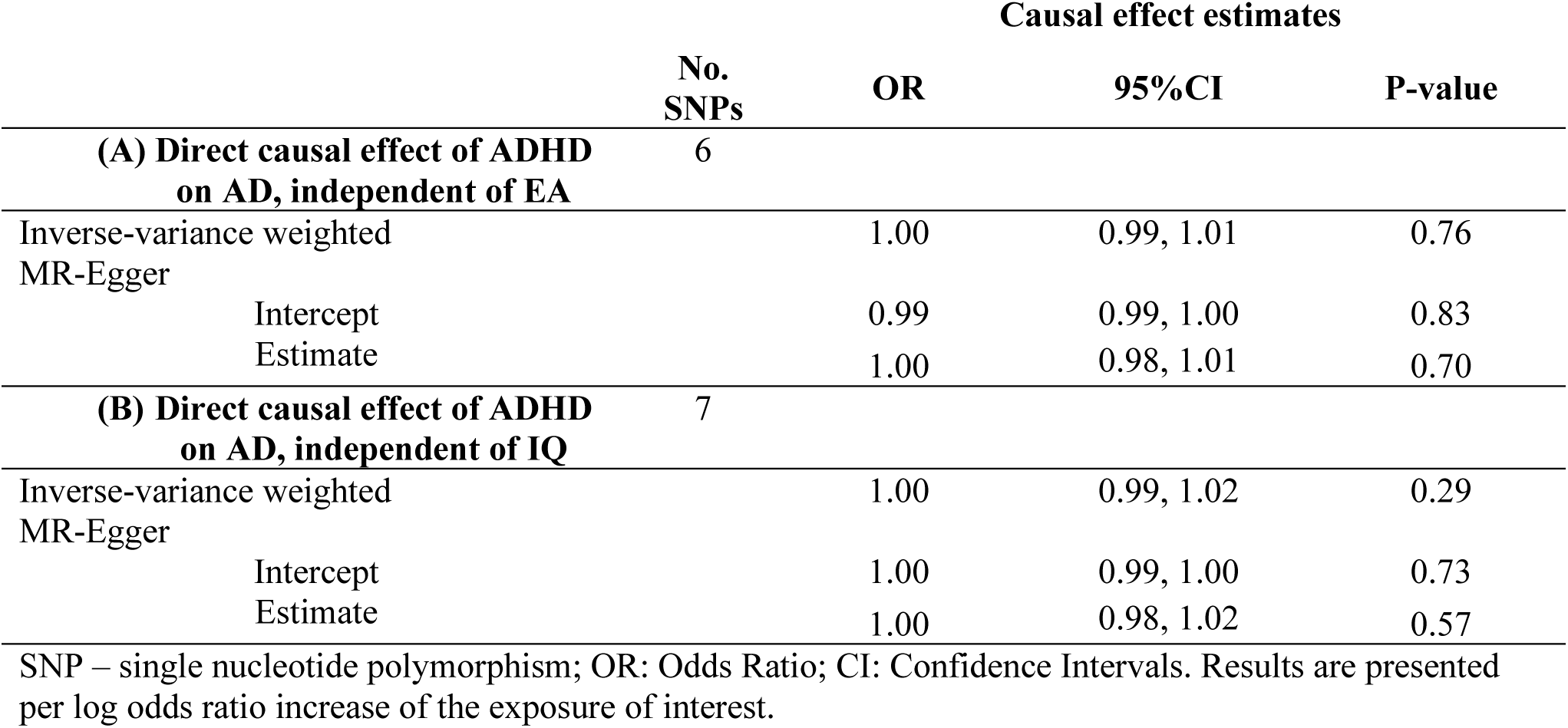
Direct causal effects of genetic liability to ADHD and on Alzheimer’s disease, independent of **(A)** Educational Attainment (**B)** Intelligence (IQ)

The results were comparable – in univariable and multivariable MR analyses - to the ones obtained from the MR analyses using Phase 1 of the Alzheimer’s disease GWAS which did not include proxy cases (Table S4-S5).

### Causal effects of genetic liability to ASD on Alzheimer’s

#### i. Bidirectional total causal effects

A total of 9 genetic variants were used to estimate the total causal effect of genetic liability to ASD on Alzheimer’s disease. We found very little evidence for a causal effect of genetic liability to ASD (OR=0.99, 95% CI: 0.97 to 1.01, p=0.70) on Alzheimer’s disease. MR-Egger and weighted median estimators produced directionally comparable results to the IVW estimator (Table 4). Additionally, no indication of horizontal pleiotropy was identified by the MR-Egger intercept term (OR=1.00, 95% CI: 0.99 to 1.00, p=0.58). No considerable heterogeneity was observed between the genetic variants (Q=6.07, p=0.63) and leave-one out analyses did not identify any SNP as influential (Figure S3A).

**Table 4.** Bidirectional effects of genetic liability to ASD on Alzheimer’s disease

|  |  | Causal effect estimates |  |  |  |
| --- | --- | --- | --- | --- | --- |
|  | No. SNPs | OR | 95%CI | P-value | Q p-value |
| ASD on Alzheimer's disease |  |  |  |  |  |
| IVW | 9 | 0.99 | 0.97,1.01 | 0.70 | 0.63 |
| MR-Egger |  |  |  |  |  |
| Intercept | 9 | 1.00 | 0.99, 1.00 | 0.58 | - |
| Slope | 9 | 0.97 | 0.89, 1.05 | 0.53 | 0.53 |
| Weighted Median | 9 | 0.99 | 0.96,1.02 | 0.66 | - |
| Alzheimer's disease on ASD |  |  |  |  |  |
| IVW | 32 | 1.19 | 0.94, 1.51 | 0.14 | 0.22 |
| MR-Egger |  |  |  |  |  |
| Intercept | 32 | 1.00 | 0.99, 1.01 | 0.53 | - |
| Slope | 32 | 1.10 | 0.77, 1.56 | 0.57 | 0.20 |
| Weighted Median | 32 | 1.17 | 0.85,1.61 | 0.31 | - |

IVW: Inverse-variance weighted; SNP – single nucleotide polymorphism; OR: Odds Ratio; CI: Confidence Intervals. Results are presented per log odds ratio increase of the exposure of interest.

In the opposite direction, a total of 32 genetic variants were used to estimate the total causal effect. We observed weak evidence that genetic liability to Alzheimer’s was associated with a higher risk of ASD (OR=1.19, 95% CI: 0.94 to 1.51, p=0.14). MR-Egger and weighted median estimators were directionally consistent with the IVW for the genetic liability of Alzheimer’s on ASD (Table 4). We had no indication that horizontal pleiotropy influenced our results as indicated by the MR-Egger intercept term (OR=1.10, 95% CI: 0.77 to 1.56, p=0.57) and by Cochrane’s Q statistic (Q=36.65, p=0.22). Moreover, leave-one out analysis did not point out any genetic variant as influential (Figure S3B).

#### ii. Direct effects

A total of 662 SNPs and 185 variants were available for inclusion in the ASD-EA-AD and ASD-IQ-AD multivariable analyses respectively. Genetic liability to ASD was not found to have direct effects on Alzheimer’s disease when educational attainment was entered in the models (OR=0.99, 95% CI: 0.98 to 1.004, p=0.28) (Table 5A). In addition, there was limited evidence to suggest a direct, independent of IQ, causal effect of genetic liability to ASD on Alzheimer’s (OR=0.99, 95%CI: 0.98 to 1.01, p=0.99) (Table 5B).

**Table 5.** Direct causal effects of genetic liability to ASD on Alzheimer’s disease, independent of **(A)** Educational Attainment (EA) **(B)** Intelligence (IQ)

|  |  | Causal effect estimates |  |  |
| --- | --- | --- | --- | --- |
|  | No.<br>SNPs | OR | 95%CI | P-value |
| (A) Direct causal effect of ASD on AD, independent of EA | 7 |  |  |  |
| Inverse-variance weighted |  | 0.99 | 0.98, 1.00 | 0.28 |
| MR-Egger |  |  |  |  |
| Intercept |  | 0.99 | 0.99, 1.00 | 0.60 |
| Estimate |  | 0.99 | 0.98, 1.01 | 0.72 |
| (B) Direct causal effect of ASD on AD, independent of IQ | 7 |  |  |  |
| Inverse-variance weighted |  | 0.99 | 0.98, 1.01 | 0.99 |
| MR-Egger |  |  |  |  |
| Intercept |  | 1.00 | 0.99, 1.00 | 0.13 |
| Estimate |  | 0.98 | 0.96, 1.00 | 0.26 |

SNP – single nucleotide polymorphism; OR: Odds Ratio; CI: Confidence Intervals. Results are presented per log odds ratio increase of the exposure of interest.

The results were comparable to the ones obtained from the MR analyses using Phase 1 of the Alzheimer’s disease GWAS which did not include proxy cases (Table S6-S7).

## DISCUSSION

Within a two-sample Mendelian randomization framework, we found limited evidence of causal effects between genetic liability to ADH and ASD on Alzheimer’s. To exclude the possibility that any causal effects might be masked by educational attainment or IQ, we conducted MVMR to estimate the direct effects of each phenotype on risk of Alzheimer’s. There was limited evidence to suggest direct causal effects of genetic liability to ADHD and ASD on Alzheimer’s risk. Finally, genetic liability to Alzheimer’s was not found to be associated with risk of ADHD and ASD.

Despite current hypotheses stemming out from observational studies that neurocognitive deficits characterizing ADHD could be associated with increased risk for Alzheimer’s disease, in the present study there was limited evidence to support this. Previous genetic studies investigating the associations between polygenic risk for Alzheimer’s and neurocognitive deficits, IQ, as well as brain structural abnormalities in childhood, have found limited evidence of associations with a genome-wide significant polygenic risk score for Alzheimer’s (36, 37). Most of the associations were identified with a polygenic risk score of Alzheimer’s using a liberal p-value threshold, suggesting possible pleiotropic pathways instead of childhood neurocognitive functioning as an early manifestation of Alzheimer’s risk (36).

However, ADHD is not a phenotypically uniform condition. Specifically, the affected children present with variable neurocognitive profiles (38), in some cases manifestations of ADHD will remit in adulthood (39, 40) and in some cases ADHD will first manifest later in life and not necessarily in childhood (late-onset ADHD) (41, 42). Therefore, different manifestations of ADHD might present different links to Alzheimer’s disease. The ADHD GWAS study we used, included a broad range of children and adults with ADHD and did not allow for testing any possible differential causal associations of genetic liability to ADHD sub-phenotypes with Alzheimer’s.

In the case of ASD, despite the hypothesis that the hyper plasticity of the ASD brain and the strong positive genetic correlations with IQ and educational attainment would reveal a protective effect against Alzheimer’s, we found limited evidence to support this. This could be attributed to the heterogeneity characterising the spectrum (43, 44), as in the case of ADHD, and possible differential causal associations across the different dimensions of the spectrum with Alzheimer’s could be speculated. Another consideration in the present study is that we included instruments at a relaxed p-value threshold of p <5×10^−07^, as only five genome-wide significant hits were identified in the ASD GWAS. This might have led to the inclusion of weak instruments, biasing the causal effect estimates towards the null (31). However, the sensitivity analyses to test for the different MR assumptions we conducted and the estimation of the strength of the included instruments, suggest that this possibility is unlikely.

Regarding both the ADHD and ASD findings of the present study, an important consideration is the possible influence of survival bias. ADHD seems to be associated with increased risk of mortality compared to the general population, which seems to arise from engagement in high risk behaviours, suicide and psychiatric comorbidity (45-47). Similarly, medication-related side effects, chronic health conditions and intellectual disabilities seem to be associated with increased mortality risk in the case of ASD (48-52). Therefore, it is possible that excess mortality associated with ADHD and ASD might bias any associations between the two conditions and Alzheimer’s.

### Strengths and Limitations

The present study is the first investigating the possible causal associations between genetic liability to ADHD, ASD and Alzheimer’s disease within a Mendelian randomization framework. This method allowed us to investigate causal relationships in the largest samples to date for the three phenotypes of interest, without the presence of confounding. Additionally, we scrutinised the validity of our findings through sensitivity analyses as well as Multivariable MR that allowed us to assess whether any causal effects were masked by IQ and educational attainment.

However, there are limitations that should be considered. The common variants that have been currently identified in the ADHD and ASD GWAS and we used as instruments, explain only a small proportion of the genetic variance of the phenotypes (14, 19). More importantly, rare variation seems to play an important part in the aetiopathogenesis of both ADHD and ASD (51, 52). We could not investigate the possible impact of rare variants associated with ADHD and ASD on risk of Alzheimer’s within the context of MR.

Also, we could not assess the possibility our MVMR results being biased due to possible sample overlap in the education attainment and Alzheimer’s GWASs, as both include samples from UK Biobank and TwinGene. However, results remained virtually the same, when we used Phase 1 of the Alzheimer’s disease GWAS which did not include proxy cases and thus had an inconsiderable overlap (1%).

### Future Directions

Investigating the possible associations of ADHD and ASD with Alzheimer’s disease is an important area of research that can have important implications for the affected individuals, their families and policy makers. It is important therefore, for future GWAS studies to offer subtype specific data in order to explore any possible differential associations of ADHD and ASD subtypes with Alzheimer’s. In addition, although longitudinal designs for this question could be difficult, the availability of registry data (e.g. Swedish registry data) can offer a unique opportunity to investigate the associations between ADHD, ASD and Alzheimer’s disease as well as the possible role of educational attainment through their extensive clinical and academic records.

## CONCLUSIONS

This is the first study to investigate the possible causal associations between genetic liability to ADHD, ASD and Alzheimer’s disease using the largest sample sizes available for each phenotype, within a Mendelian randomization framework. We found limited evidence to suggest total and direct effects of genetic liability to ADHD and ASD on risk of Alzheimer’s disease. We hope that this will be an important step towards encouraging future research into possible differential associations of ADHD and ASD subtypes and risk of Alzheimer’s disease as well as utilizing longitudinal data.

## Notes

**Funding:** This work was supported by a grant from the BRACE Alzheimer’s charity (BR16/028). PP, BL, RKL, ELA, GDS, LDH and ES work in a unit that receives funding from the University of Bristol and the UK Medical Research Council (MC_UU_00011/1, MC_UU_00011/3 and MC_UU_00011/6). CD is funded by the Wellcome Trust (grant ref: 108902/B/15/Z). BL was supported by a research grant from the Wellcome Trust (grant ref: 204895/Z/16/Z). RKL is supported by a Wellcome Trust PhD studentship (Grant ref: 215193/Z18/Z). LDH and ELA are funded by a Career Development Award from the UK Medical Research Council (MR/M020894/1 and MR/P014437/1, respectively). This publication is the work of the authors, and ES will serve as a guarantor for the contents of this paper.

**Competing interests:** None declared

### Competing Interest Statement

The authors have declared no competing interest.

