## Supplementary Material for "Exploring the causal effects of genetic liability to ADHD and Autism on Alzheimer’s disease"

**TABLES**

### A. Main analyses supplementary material

**Supplementary Table S1.** Characteristics of 10 genetic variants associated with genetic liability to ADHD

**Supplementary Table S2.** Characteristics of 9 genetic variants associated with genetic liability to ASD

**Supplementary Table S3.** Characteristics of 32 genetic variants associated with genetic liability to Alzheimer's disease

### B. Additional analyses results

**Supplementary Table S4.** Bidirectional effects of genetic liability to ADHD on Alzheimer's disease, using Phase 1 of the Alzheimer's disease GWAS which did not include proxy cases

**Supplementary Table S5.** Direct causal effects of genetic liability to ADHD and on Alzheimer's disease, independent of **(A)** Educational Attainment **(B)** Intelligence (IQ), using Phase 1 of the Alzheimer's disease GWAS which did not include proxy cases

**Supplementary Table S6.** Bidirectional effects of genetic liability to ASD on Alzheimer's disease, using Phase 1 of the Alzheimer's disease GWAS which did not include proxy cases

**Supplementary Table S7.** Direct causal effects of genetic liability to ASD and on Alzheimer's disease, independent of **(A)** Educational Attainment **(B)** Intelligence (IQ), using Phase 1 of the Alzheimer's disease GWAS which did not include proxy cases

### A. Main analyses supplementary material

**Table S1.** Characteristics of 10 genetic variants associated with genetic liability to ADHD

| SNP ID | Chr: pos | Locus | Effect/ Ref allele | ADHD | | AD | |
|---|---|---|---|---|---|---|---|
| | | | | Beta (SE) | P-value | Beta (SE) | P-value |
| rs11420276 | 1:43718521 | LOC101929592, ST3GAL3 | G/GT | 0.1071 (0.0149) | 6.45E-13 | -0.0028 (0.0117) | 8.05E-01 |
| rs1222063 | 1:96136884 | - | A/G | 0.0962 (0.0174) | 3.07E-08 | -0.0074 (0.0110) | 5.01E-01 |
| rs4858241 | 3:20669071 | Intergenic | T/G | 0.0821 (0.0143) | 8.17E-09 | -0.0004 (0.0022) | 8.22E-01 |
| rs28411770 | 4:31149834 | - | T/C | 0.0861 (0.0151) | 1.15E-08 | -0.0102 (0.0110) | 3.52E-01 |
| rs4916723 | 5:87854395 | LINC00461, MIR9-2, LINC02060, TMEM161B-AS1 | C/A | 0.0777 (0.0138) | 1.81E-08 | -0.0016 (0.0023) | 4.69E-01 |
| rs10262192 | 7:114451698 | FOXP2 | A/G | 0.0740 (0.0135) | 3.66E-08 | 0.0026 (0.0021) | 2.22E-01 |
| rs74760947 | 8:34352610 | LINC01288 | G/A | 0.1796 (0.0317) | 1.39E-08 | 0.0064 (0.0053) | 2.25E-01 |
| rs1427829 | 12:89760744 | DUSP6, POC1B | A/G | 0.0821 (0.0136) | 1.35E-09 | 0.0023 (0.0021) | 2.74E-01 |
| rs8039398 | 15:47438673 | SEMA6D | C/T | 0.0799 (0.0135) | 2.99E-09 | 0.0015 (0.0021) | 4.51E-01 |
| rs212178 | 16:72578131 | LINC01572 | G/A | 0.1170 (0.0205) | 1.20E-08 | -0.0025 (0.0034) | 4.56E-01 |

**Table S2.** Characteristics of 9 genetic variants associated with genetic liability to ASD

| SNP ID | Chr: pos | Locus | Effect/ Ref allele | ASD | | AD | |
|---|---|---|---|---|---|---|---|
| | | | | Beta (SE) | P-value | Beta (SE) | P-value |
| rs2391769 | 1:96513405 | LOC105378866 | G/A | 0.0769 (0.0145) | 1.14E-07 | 0.0001 (0.0022) | 9.64E-01 |
| rs6701243 | 1:98627228 | - | A/C | 0.0735 (0.0144) | 3.07E-07 | -0.0004 (0.0023) | 8.57E-01 |
| rs1452075 | 3:62495388 | CADPS | T/C | 0.0807 (0.0155) | 2.07E-07 | 0.0006 (0.0023) | 7.65E-01 |
| rs325485 | 5:104659667 | LOC105379109 | A/G | 0.0728 (0.0143) | 3.25E-07 | -0.0008 (0.0021) | 6.82E-01 |
| rs111931861 | 7:105103772 | KMT2E | G/A | 0.2169 (0.0409) | 1.12E-07 | 0.0276 (0.0231) | 2.33E-01 |
| rs45595836 | 10:16649400 | RSU1 | T/C | 0.1389 (0.0272) | 3.13E-07 | -0.0008 (0.0040) | 8.34E-01 |
| rs112635299 | 14:94371805 | - | T/G | 0.2209 (0.0432) | 3.04E-07 | -0.0103 (0.0089) | 2.48E-01 |
| rs11481126 | 20:14841527 | MACROD2 | GA/G | 0.0737 (0.0140) | 1.26E-07 | 0.0185 (0.0107) | 8.34E-02 |

| | | | | | | | |
|---|---|---|---|---|---|---|---|
| rs910805 | 20:21267478 | - | G/A | 0.0956 (0.0160) | 2.04E-09 | -0.0008 (0.0025) | 7.30E-01 |

**Table S3.** Characteristics of 32 genetic variants associated with genetic liability to Alzheimer's disease

| SNP ID | Chr: pos | Locus | Effect/ Ref allele | AD | | ADHD | | ASD | |
|---|---|---|---|---|---|---|---|---|---|
| | | | | Beta (SE) | P-value | Beta (SE) | P-value | Beta (SE) | P-value |
| rs2093760 | 1:207613483 | CR1 | A/G | 0.0235 (0.0026) | 1.10E-18 | 0.0065 (0.017) | 6.98E-01 | 0.0218 (0.0175) | 2.10E-01 |
| rs4575098 | 1:161185602 | B4GALT3 | A/G | 0.0159 (0.0025) | 2.05E-10 | -0.0306 (0.0161) | 5.75E-02 | -0.0059 (0.0165) | 7.16E-01 |
| rs4663105 | 2:12713385 | LOC105373605 | C/A | 0.0302 (0.0021) | 3.38E-44 | 0.0151 (0.0145) | 2.99E-01 | 0.0095 (0.0143) | 5.04E-01 |
| rs6448453 | 4:11024404 | - | A/G | 0.0143 (0.0023) | 1.93E-09 | -0.0227 (0.0154) | 1.38E-01 | -0.0133 (0.0157) | 3.93E-01 |
| rs143332484 | 6:41161469 | TREM2 | T/C | 0.0716 (0.0129) | 3.01E-08 | -0.0031 (0.0674) | 9.62E-01 | -0.0652 (0.0645) | 3.12E-01 |
| rs9381563 | 6:47464901 | - | C/T | 0.0140 (0.0022) | 2.52E-10 | 0.0051 (0.0148) | 7.31E-01 | 0.0108 (0.0146) | 4.57E-01 |
| rs1859788 | 7:100374211 | PILRA | G/A | 0.0178 (0.0022) | 2.22E-15 | 0.0263 (0.0147) | 7.25E-02 | 0.0033 (0.0150) | 8.20E-01 |
| rs7810606 | 7:143411065 | EPHA1-AS1 | C/T | 0.0140 (0.0021) | 3.59E-11 | 0.0282 (0.0143) | 4.74E-02 | 0.0303 (0.0143) | 3.38E-02 |
| rs28834970 | 8:27337604 | PTK2B | C/T | 0.0148 (0.0021) | 1.04E-11 | 0.0043 (0.014) | 7.54E-01 | 0.0262 (0.0143) | 6.75E-02 |
| rs4236673 | 8:27607412 | CLU | G/A | 0.0195 (0.0021) | 2.61E-19 | -0.0266 (0.0146) | 6.86E-02 | -0.0121 (0.0142) | 3.90E-01 |
| rs11257238 | 10:11675398 | LOC105376412, LOC105376413 | C/T | 0.0125 (0.0022) | 1.26E-08 | 0.01159 (0.0142) | 4.13E-01 | 0.0083 (0.0143) | 5.54E-01 |
| rs11218343 | 11:121564878 | SORL1 | T/C | 0.0348 (0.0051) | 1.09E-11 | 0.0212 (0.0355) | 5.50E-01 | 0.0419 (0.0363) | 2.48E-01 |
| rs2081545 | 11:60190907 | - | C/A | 0.0173 (0.0021) | 1.55E-15 | -0.0240 (0.0138) | 8.07E-02 | -0.0182 (0.0141) | 1.96E-01 |
| rs867611 | 11:86065502 | PICALM | A/G | 0.0198 (0.0022) | 2.19E-18 | 0.01230 (0.0143) | 3.93E-01 | -0.0146 (0.0147) | 3.20E-01 |
| rs12590654 | 14:92472511 | SLC24A4 | G/A | 0.0143 (0.0022) | 1.65E-10 | -0.0018 (0.0149) | 9.00E-01 | 0.0039 (0.0149) | 7.90E-01 |
| rs442495 | 15:58730416 | ADAM10 | T/C | 0.0133 (0.0022) | 1.31E-09 | 0.01460 (0.0144) | 3.08E-01 | -0.0053 (0.0147) | 7.16E-01 |
| rs117618017 | 15:63277703 | APH1B | T/C | 0.0175 (0.0031) | 3.35E-08 | **NA** | **NA** | 0.0200 (0.0227) | 3.75E-01 |
| rs59735493 | 16:31121779 | KAT8 | G/A | 0.0126 (0.0023) | 3.98E-08 | -0.0162 (0.0153) | 2.85E-01 | -0.0016 (0.0153) | 9.09E-01 |
| rs28394864 | 17:49373413 | LOC102724596 | A/G | 0.0119 (0.0021) | 1.87E-08 | 0.0344 (0.0136) | 1.15E-02 | 0.0352 (0.0139) | 1.15E-02 |
| rs113260531 | 17:5235685 | SCIMP, LOC100130950 | A/G | 0.0194 (0.0031) | 9.16E-10 | -0.0055 (0.0209) | 7.90E-01 | -0.0411 (0.0215) | 5.60E-02 |
| rs10422568 | 19:44695207 | LOC107985305 | C/T | 0.0336 (0.0053) | 2.94E-10 | 0.0442 (0.0358) | 2.17E-01 | 0.0598 (0.0364) | 9.97E-02 |
| rs2965169 | 19:44747899 | BCL3 | A/C | 0.0340 (0.0021) | 3.13E-57 | 0.0202 (0.0159) | 2.05E-01 | 0.0162 (0.0149) | 2.73E-01 |
| rs2967669 | 19:44798685 | CBLC | A/G | 0.0306 (0.0030) | 2.11E-23 | 0.0111 (0.0251) | 6.56E-01 | -0.0061 (0.0223) | 7.82E-01 |
| rs12978931 | 19:44860443 | NECTIN2 | A/G | 0.0313 (0.0026) | 1.49E-32 | 0.0106 (0.0180) | 5.57E-01 | 0.0172 (0.0181) | 3.42E-01 |
| rs138607350 | 19:44860563 | NECTIN2 | G/T | 0.2187 (0.0128) | 2.94E-65 | 0.0208 (0.0673) | 7.58E-01 | 0.0654 (0.0632) | 3.01E-01 |
| rs77301115 | 19:44893716 | TOMM40 | A/G | 0.2065 (0.0067) | 1.00E-200 | 0.0247 (0.0418) | 5.54E-01 | 0.0222 (0.0423) | 5.99E-01 |
| rs204907 | 19:44958739 | CLPTM1 | A/G | 0.0303 (0.0046) | 9.04E-11 | -0.0242 (0.0350) | 4.90E-01 | -0.0011 (0.0357) | 9.76E-01 |

| | | | | | | | | | |
|---|---|---|---|---|---|---|---|---|---|
| rs117612135 | 19:45001269 | RELB | T/C | 0.1374 (0.0095) | 1.12E-46 | -0.0292 (0.0682) | 6.68E-01 | -0.0192 (0.0641) | 7.63E-01 |
| rs346773 | 19:45229943 | EXOC3L2 | T/C | 0.0330 (0.0034) | 3.66E-21 | -0.0319 (0.0264) | 2.25E-01 | -0.0203 (0.0258) | 4.28E-01 |
| rs123187 | 19:45327689 | - | G/A | 0.0118 (0.0021) | 3.70E-08 | -0.0259 (0.0141) | 6.44E-02 | 0.0211 (0.0143) | 1.38E-01 |
| rs3865444 | 19:51224706 | CD33 | C/A | 0.0133 (0.0022) | 6.34E-09 | 0.0011 (0.0144) | 9.41E-01 | -0.0176 (0.0145) | 2.26E-01 |
| rs6014724 | 20:56423488 | CASS4 | A/G | 0.0222 (0.0035) | 6.56E-10 | 0.0045 (0.0251) | 8.54E-01 | 0.0078 (0.0254) | 7.55E-01 |

### B. Additional analyses results

**Table S4.** Bidirectional effects of genetic liability to ADHD on Alzheimer's disease, using Phase 1 of the Alzheimer's disease GWAS which did not include proxy cases

| | | **Causal effect estimates** | | | |
|---|---|---|---|---|---|
| | **No. SNPs** | **OR** | **95%CI** | **P-value** | **Q p-value** |
| **ADHD on Alzheimer's disease** | | | | | |
| IVW | 10 | 1.01 | 0.87, 1.18 | 0.82 | 0.38 |
| MR-Egger | | | | | |
| Intercept | 10 | 0.96 | 0.90, 1.02 | 0.26 | - |
| Slope | 10 | 1.52 | 0.73, 3.17 | 0.25 | 0.40 |
| Weighted Median | 10 | 1.08 | 0.87,1.32 | 0.46 | - |
| **Alzheimer's disease on ADHD** | | | | | |
| IVW | 25 | 1.01 | 0.97, 1.05 | 0.40 | 0.63 |
| MR-Egger | | | | | |
| Intercept | 25 | 1.00 | 0.98, 1.01 | 0.90 | - |
| Slope | 25 | 1.01 | 0.94, 1.09 | 0.73 | 0.57 |
| Weighted Median | 25 | 1.00 | 0.95,1.06 | 0.74 | - |

IVW: Inverse-variance weighted; SNP – single nucleotide polymorphism; OR: Odds Ratio; CI: Confidence Intervals. Results are presented per log odds ratio increase of the exposure of interest.

**Table S5.** Direct causal effects of genetic liability to ADHD and on Alzheimer's disease, independent of (A) Educational Attainment (B) Intelligence (IQ), using Phase 1 of the Alzheimer's disease GWAS which did not include proxy cases

| | No. SNPs | Causal effect estimates | | |
|---|---|---|---|---|
| | | **OR** | **95%CI** | **P-value** |
| **(A) Direct causal effect of ADHD on AD, independent of EA** | 6 | | | |
| Inverse-variance weighted | | 1.02 | 0.95, 1.11 | 0.45 |
| MR-Egger | | | | |
| Intercept | | 0.99 | 0.99, 1.00 | 0.74 |
| Estimate | | 1.04 | 0.94, 1.15 | 0.43 |
| **(B) Direct causal effect of ADHD on AD, independent of IQ** | | | | |
| Inverse-variance weighted | | 0.99 | 0.88, 1.10 | 0.86 |
| MR-Egger | | | | |
| Intercept | | 1.00 | 0.99, 1.00 | 0.87 |
| Estimate | | 0.98 | 0.84, 1.14 | 0.81 |

SNP – single nucleotide polymorphism; OR: Odds Ratio; CI: Confidence Intervals. Results are presented per log odds ratio increase of the exposure of interest.

**Table S6.** Bidirectional effects of genetic liability to ASD on Alzheimer’s disease, using Phase 1 of the Alzheimer’s disease GWAS which did not include proxy cases

| | | Causal effect estimates | | | |
|---|---|---|---|---|---|
| | **No. SNPs** | **OR** | **95%CI** | **P-value** | **Q p-value** |
| **ASD on Alzheimer’s disease** | | | | | |
| IVW | 9 | 1.05 | 0.85, 1.29 | 0.64 | 0.17 |
| MR-Egger | | | | | |
| Intercept | 9 | 0.94 | 0.87, 1.03 | 0.23 | - |
| Slope | 9 | 1.92 | 0.68, 5.38 | 0.21 | 0.20 |
| Weighted Median | 9 | 1.00 | 0.81, 1.25 | 0.92 | - |
| **Alzheimer’s disease on ASD** | | | | | |
| IVW | 28 | 1.04 | 1.00, 1.08 | **0.01** | 0.52 |
| MR-Egger | | | | | |
| Intercept | 28 | 0.99 | 0.98, 1.00 | 0.60 | - |
| Slope | 28 | 1.06 | 0.99, 1.14 | 0.08 | 0.48 |
| Weighted Median | 28 | 1.06 | 1.00, 1.12 | **0.02** | - |

IVW: Inverse-variance weighted; SNP – single nucleotide polymorphism; OR: Odds Ratio; CI: Confidence Intervals. Results are presented per log odds ratio increase of the exposure of interest.

**Table S7.** Direct causal effects of genetic liability to ASD and on Alzheimer's disease, independent of **(A)** Educational Attainment **(B)** Intelligence (IQ), using Phase 1 of the Alzheimer's disease GWAS which did not include proxy cases

| | No. SNPs | Causal effect estimates | | |
|---|---|---|---|---|
| | | **OR** | **95%CI** | **P-value** |
| **(A) Direct causal effect of ASD on AD, independent of EA** | 7 | | | |
| Inverse-variance weighted | | 0.99 | 0.93, 1.06 | 0.98 |
| MR-Egger | | | | |
| Intercept | | 0.99 | 0.99, 1.00 | 0.80 |
| Estimate | | 1.00 | 0.91, 1.11 | 0.87 |
| **(B) Direct causal effect of ASD on AD, independent of IQ** | 7 | | | |
| Inverse-variance weighted | | 1.00 | 0.88, 1.12 | 0.98 |
| MR-Egger | | | | |
| Intercept | | 0.99 | 0.99, 1.00 | 0.92 |
| Estimate | | 1.00 | 0.84, 1.20 | 0.93 |

SNP – single nucleotide polymorphism; OR: Odds Ratio; CI: Confidence Intervals. Results are presented per log odds ratio increase of the exposure of interest.

**FIGURES**

**Figure S1.** Flow diagram illustrating the identification, extraction and harmonization process of the variants included across the analyses of the present study.

**Figure S2.** Leave-one out analyses of genetic liability to **(A)** ADHD on AD and **(B)** AD on ADHD, using IVW. Estimates shown as log odds and 95%CI

**Figure S3.** Leave-one out analyses of genetic liability to **(A)** ASD on AD and **(B)** AD on ASD, using IVW. Estimates shown as log odds and 95%CI

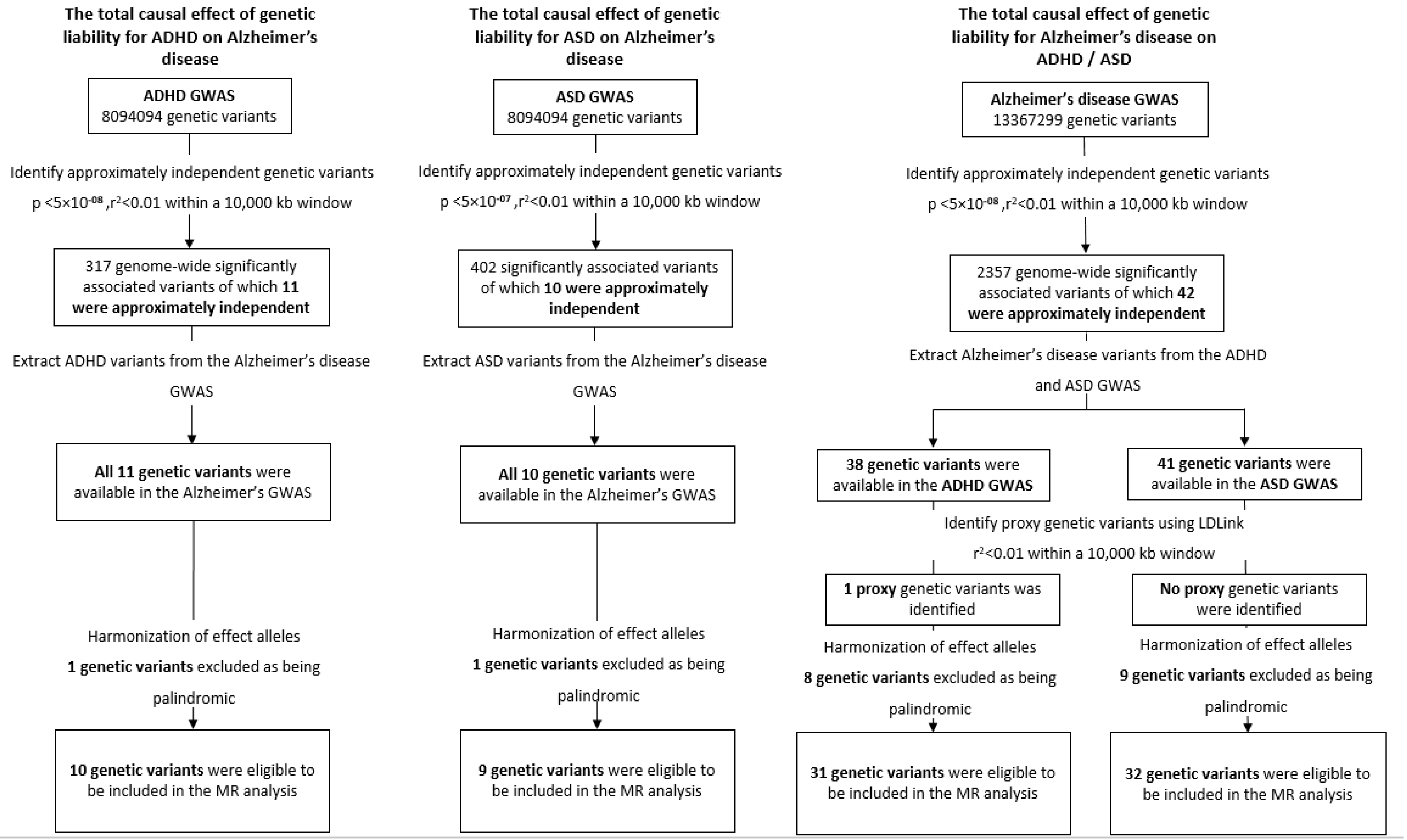


**Figure S1.** Flow diagram illustrating the identification, extraction and harmonization process of the variants included across the analyses of the present study.

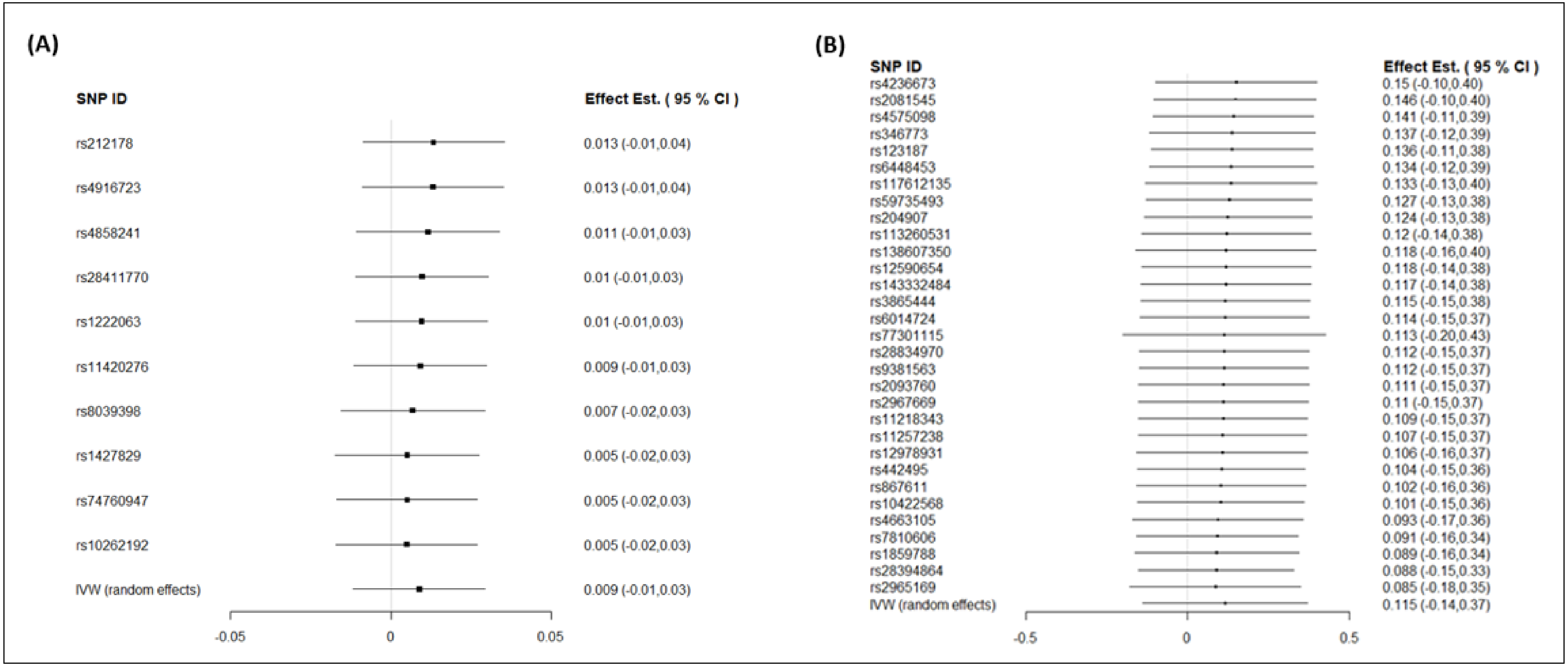


**Figure S2.** Leave-one out analyses of genetic liability to ***(A)*** ADHD on AD and ***(B)*** AD on ADHD, using IVW. Estimates shown as log odds and 95%CI

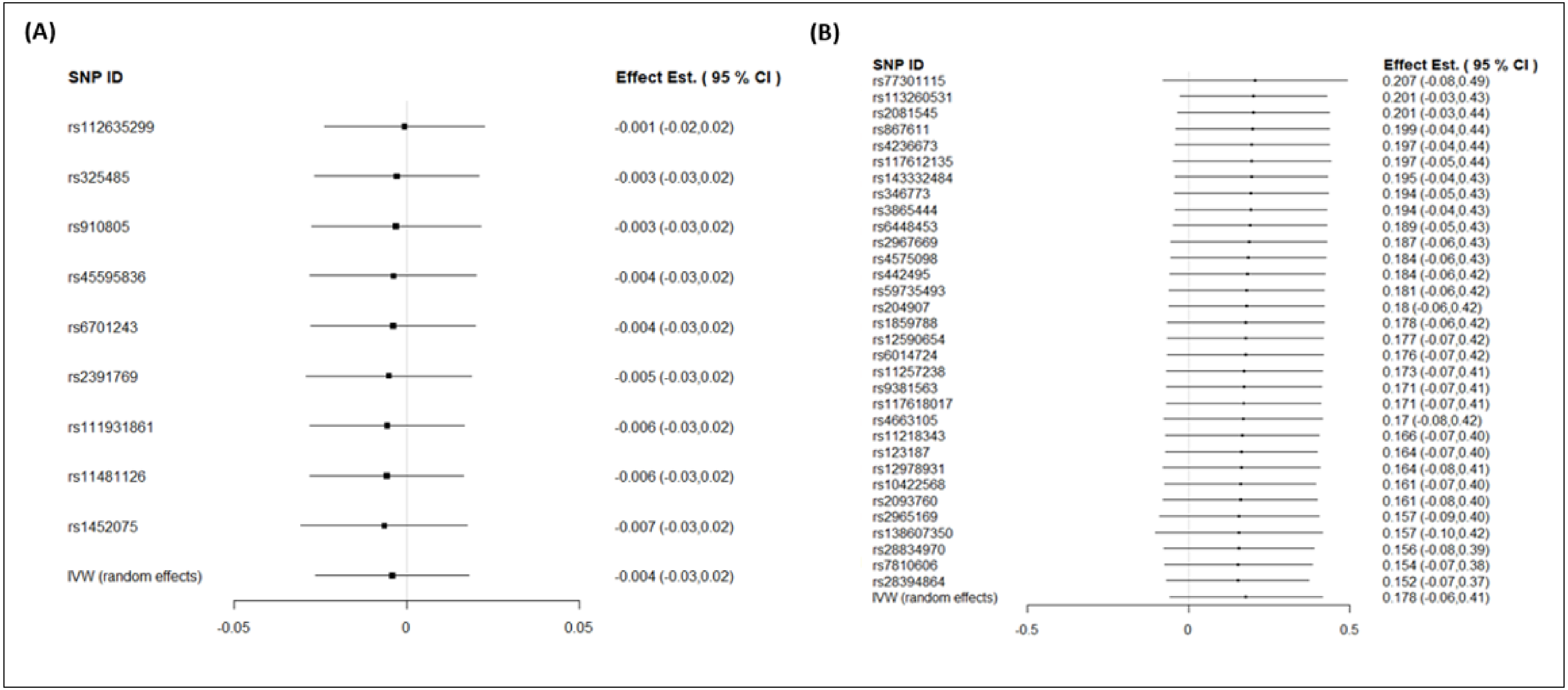


**Figure S3.** Leave-one out analyses of genetic liability to ***(A)*** ASD on AD and ***(B)*** AD on ASD, using IVW. Estimates shown as log odds and 95%CI
